# The Evolution of Insect Metallothioneins

**DOI:** 10.1101/2020.06.25.172213

**Authors:** Mei Luo, Cédric Finet, Haosu Cong, Hong-yi Wei, Henry Chung

## Abstract

Metallothioneins (MTs) are a family of cysteine-rich metal-binding proteins that are important in the chelating and detoxification of toxic heavy metals. Until now, the short length and the low sequence complexity of MTs has hindered the possibility of any phylogenetic reconstruction, hampering the study of their evolution. To answer this longstanding question, we developed an iterative BLAST search pipeline that allowed us to build a unique dataset of more than 300 MT sequences in insects. By combining phylogenetics and synteny analysis, we reconstructed the evolutionary history of MTs in insects. We show that the MT content in insects has been shaped by lineage-specific tandem duplications from a single ancestral MT. Strikingly, we also uncovered a sixth MT, *MtnF*, in the model organism *Drosophila melanogaster*. *MtnF* evolves faster than other MTs and is characterized by a non-canonical length and higher cysteine content. Our methodological framework not only paves the way for future studies on heavy metal detoxification but also can allow us to identify other previously unidentified genes and other low complexity genomic features.

## INTRODUCTION

Heavy metals like copper (Cu) and zinc (Zn) have been co-opted over time as essential components of numerous transcriptional factors and catalytic enzymes (Hamer 1986). However, high concentration of heavy metals can be cytotoxic, and organisms have evolved intricate strategies to detoxify and excrete heavy metals. Over time, these detoxification strategies have also been used to detoxify other non-essential heavy metals such as cadmium (Cd) (Klaassen *et al.* 1999). The detoxification strategies are diverse and can often be taxon-specific (Cobbett and Goldsbrough 2002; Navarro and Schneuwly 2017). One common mechanism among many organisms is the use of low molecular weight, cysteine-rich peptides known as metallothioneins (MTs) to chelate and regulate the concentration of heavy metals in the cell (Hamer 1986).

Since their seminal discovery in the horse kidney (Margoshes et al., 1957), MTs have been discovered in both eukaryotes and prokaryotes. However, because of their short sequences (~60 amino acids) and low sequence complexity due to their high cysteine content, it has been claimed that phylogenetic analysis cannot be performed (Binz and Kägi 1999; Ziller and Fraissinet-Tachet 2018). Instead, MTs have been classified into 15 families according to the organism they are isolated from (Binz and Kägi 1999) or to their function as Zn- or Cu-thioneins (Palacios *et al.* 2011). While most organisms have Cu-thioneins, Zn-thioneins have only been found in higher organisms such as the Metazoa (Capdevila and Atrian 2011; Wang *et al.* 2014). Another reason that makes phylogenetic analyses of MTs difficult is the scarcity of annotated MT sequences in sequenced genomes. Indeed, *in silico* gene predictions often fail to identify MT-encoding genes that contain very small exons punctuated by large introns (Ragusa *et al.* 2017; Purać *et al.* 2019).

MTs have been intensively studied in the fruit fly *Drosophila melanogaster*. *MtnA* was the first member of this gene family cloned from copper-fed larvae (Lastowski-Perry *et al.* 1985), followed by *MtnB* cloned from a cadmium-resistant *Drosophila* cell line (Mokdad *et al.* 1987). More than a decade later, the release of the *D. melanogaster* genome (Adams *et al.* 2000) allowed the identification of *MtnC* and *MtnD* by sequence similarity (Egli *et al.* 2003). The four *MtnA-D* genes are inducible by copper and cadmium through the binding of the transcription factor MTF-1 (Egli *et al.* 2003). Further work showed that *MtnA* knockout flies are sensitive to copper, *MtnB* knockout flies are sensitive to cadmium, whereas *MtnC* and *MtnD* knockout flies do not show any differences in copper or cadmium resistance, suggesting that these MTs have distinct roles in heavy metal detoxification (Egli *et al.* 2006). In 2011, a fifth member, *MtnE*, was discovered in the *D. melanogaster* genome through bioinformatic analysis (Atanesyan *et al.* 2011). *MtnE* is also inducible by heavy metals such as copper (Atanesyan *et al.* 2011), and it is classified as a Cu-thionein like the other four *Drosophila* MTs (Pérez-Rafael *et al.* 2012; Navarro and Schneuwly 2017). A striking feature is the absence of Zn-thioneins in *D. melanogaster*, and more generally in insects, whereas they are found in all metazoans (Atrian 2009). Outside the *Drosophila* genus (Guirola *et al.* 2011), only 14 MT genes have been published in insects (Suppl Table 1). There is only one MT in the honey bee *Apis mellifera* (Purać *et al.* 2019), two MTs in the Chinese grasshopper *Oxya chinensis* (Liu *et al.* 2014), but five MTs in sequenced *Drosophila* species genomes (Guirola *et al.* 2011). The MT content is dynamic and evolves rapidly across insects. Furthermore, insects live in diverse environments with different heavy metal challenges (Janssens *et al.* 2009; Merritt and Bewick 2017; Navarro and Schneuwly 2017), offering a good model to study how heavy metal detoxification evolves.

To investigate how insect MTs evolve, we built a comprehensive dataset of MT sequences encompassing the main insect orders. We took benefit from the recent release of a huge amount of genomic and transcriptomic data in insects (Yin *et al.* 2015; Thomas *et al.* 2020). To avoid dealing with large introns in genomes, we developed a iterative BLAST search pipeline on all available insect transcriptomes (Yin *et al.* 2015), before using this dataset to further identify MTs in sequenced insect genomes. In total, we identified and annotated more than 300 insect MTs from about 100 insect species based on available sequenced genomes and transcriptomes. Using a combination of phylogenetic and synteny analyses, we showed that the insect MTs evolved from one single ancestral MT gene in the last common ancestor after the diversification of insects. We have also discovered *MtnF*, a putative sixth metallothionein in the Diptera, including *D. melanogaster* in spite of previous intensive work in this model species. MtnF reveals non-canonical features compared to other insect MTs, suggesting putative different binding specificities.

## RESULTS AND DISCUSSION

### Combined phylogenetic and synteny analyses reveal a single ancestral MT in insects

The study of MT family evolution is essential to understanding the molecular response to heavy metal and its diversification. However, our knowledge of MT evolution has been hampered by technical difficulties in identifying MT sequences in genomes and transcriptomes. Indeed, MT genes contain large introns, and MT sequences are short (~120-200 bp) and highly divergent between genera. To facilitate comparative studies of MTs in insects, we reasoned that a large sampling should be of help. We applied an exhaustive BLAST search to around a hundred insect available transcriptomes. We identified more than 300 MT sequences encompassing 13 orders across different insect species (**Table 1**), whereas the number of published MTs was 19 prior to this study (**Table S1**).

**Table 1.** Summary of MT content in sampled insect species.

| Order | number of sampled species | number of MTs |
| --- | --- | --- |
| Blattodea | 2 | 3 |
| Coleoptera | 11 | 13 |
| Dermaptera | 1 | 2 |
| Diptera | 46 | 195 |
| Hemiptera | 7 | 8 |
| Hymenoptera | 31 | 35 |
| Lepidoptera | 27 | 73 |
| Neuroptera | 1 | 2 |
| Odonata | 1 | 2 |
| Orthoptera | 2 | 3 |
| Phasmatodea | 4 | 6 |
| Plecoptera | 1 | 3 |
| Siphonaptera | 1 | 1 |
| Total | 135 | 346 |

We performed maximum-likelihood phylogenetic reconstruction on a 202-taxon MT dataset. The tree obtained clarifies the number of main MT lineages in insects. In this tree, the coleopteran MTs are clustered in a single clade, like the hymenopteran MTs, suggesting they derive from a single MT in the last common ancestor of the extant Coleoptera and Hymenoptera, respectively. The lepidopteran MTs split into three main clades that we named clades α, β and γ (**Figure 1A**). This result suggests that at least three MT genes were present in the last common ancestor of the extant Lepidoptera. Similarly, the dipteran MTs split into several clades, suggesting multiple ancestral MTs in the Diptera. The obtained tree also showed that the lepidopteran clade γ groups with dipteran clades (**Figure 1A**). The branch leading to this Lepidoptera-Diptera superclade is well supported with a bootstrap value of 73 (**Figure 1A**). Because Lepidoptera and Diptera are two closely-related orders (Misof *et al.* 2014), our data support a MT gene duplication prior to the divergence between Lepidoptera and Diptera.

**Figure 1.**
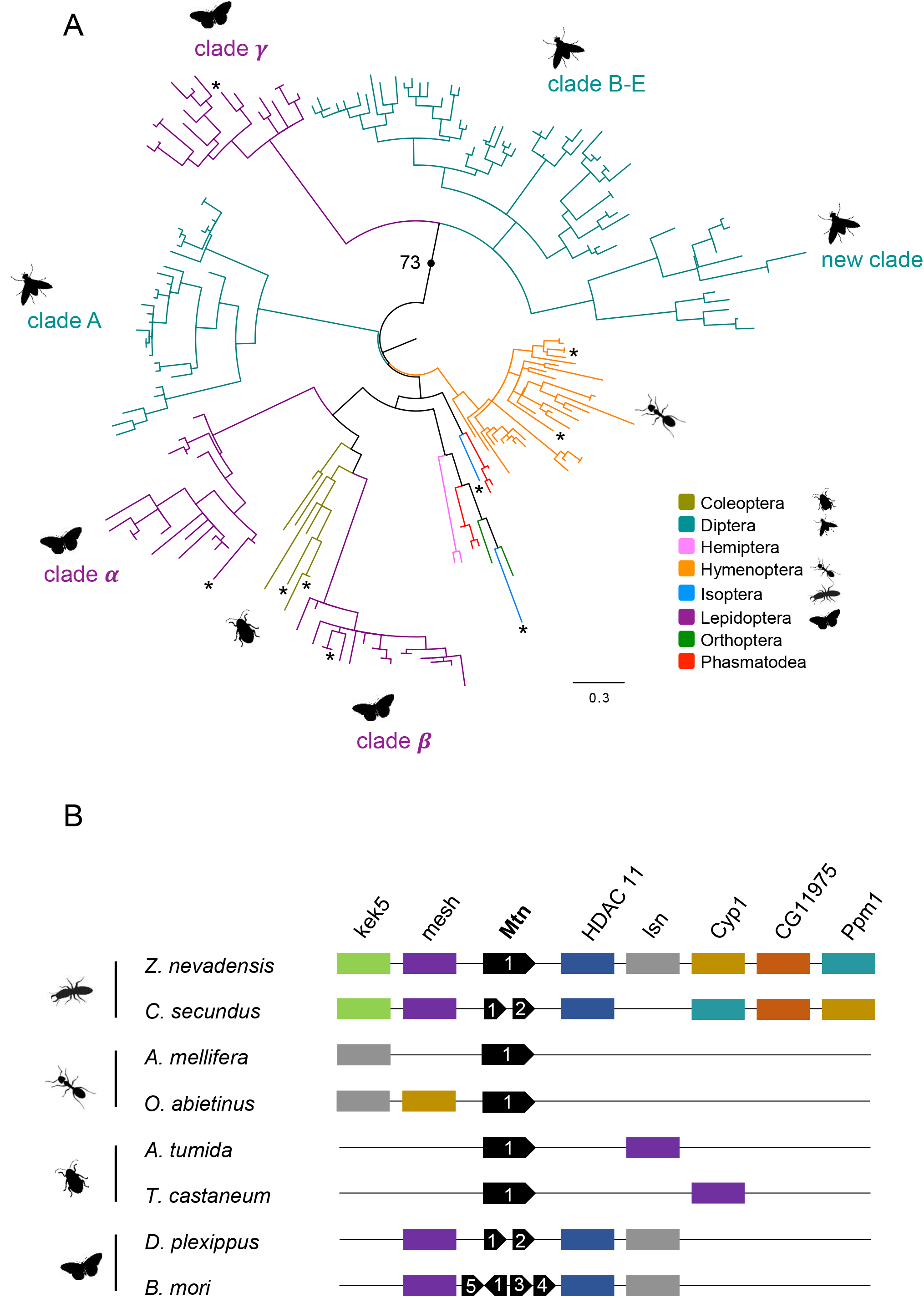
Phylogeny and synteny of MT genes in insects. (A) Phylogram of the 202-taxon analyses obtained through PhyML maximum-likelihood reconstruction. Analyses were conducted using the LG+Γ model. Support value is shown for selected branches (all branch supports are shown in online supplementary information). Scale bar indicates number of changes per site. (B) The conservation of synteny was investigated in taxa with available genomes (*), suggesting a single MT locus in the last common ancestor of extant insects.

Due to their short size and high divergence, MTs are however limiting phylogenetic markers. To provide further evidence for our aforementioned observations, we also investigated the conservation of synteny at MT loci as an independent and supplementary approach (Finet *et al.* 2019; Vakirlis *et al.* 2020). Taxa for which genomic sequences were available are indicated by an asterisk onto the phylogenetic tree (**Figure 1A**). We found that the MT sequences are flanked by the same genes in the Isoptera (*Zootermopsis nevadensis* and *Cryptotermes secundus*), in the Hymenoptera (*Apis mellifera* and *Orussus abietinus*), in the Coleoptera (*Aethina tumida* and *Tribolium castaneum*), and in the Lepidoptera (*Danaus plexippus* and *Bombyx mori*) (**Figure 1B**). Along with the presence of orthologues between Lepidoptera and Diptera, this deep conservation of microsynteny across the insects suggests the existence of a single MT gene in the last common ancestor of extant insects.

It is noteworthy that dipteran MT genes are not located at the same genomic locus shared with other insects. Nevertheless, the shared flanking genes *mesh*, *HDAC11*, *lsn*, and *CG11975* have been identified on the chromosome 3R in *D. melanogaster*, where the MTs are located. The high rate of chromosomal rearrangement in the Diptera (Ranz *et al.* 2001; Adler *et al.* 2016; Artemov *et al.* 2017; Stewart and Rogers 2019) might explain the breakdown of the ancestral MT locus in the Diptera. In particular, the genomic locus surrounding *MtnF* is deeply rearranged in *D. buzzatii* (Calvete *et al.* 2012). On the contrary, the conservation of microsynteny at the MT locus in the Lepidoptera is in line with low rates of change in gene order that have been found in this order based on coarse-scale mapping data (Pringle *et al.* 2007; d’Alencon *et al.* 2010). Whereas conservation of microsynteny between Hemiptera and other insects has been reported (Finet *et al.* 2018; Mandrioli *et al.* 2019), we did not find clear evidence of microsynteny at the MT locus in Hemiptera. Future increase of the current scarce genomic data in Hemiptera should help to conclude whether the MT locus has been translocated.

### Insect MT repertoire originated through tandem duplication

A widespread feature of the insect MT gene family is the presence of several copies in tandem in sequenced insect genomes. For example, the moth *B. mori* has four MT genes in tandem (**Figure 1B**) and the housefly *Musca domestica* has four clustered copies of *MntA* (**Figure 2A)**. This is also true for earliest-diverging lineages of insects like *C. secundus* for which we identified two MT copies in tandem (**Figure 1B**). Similar results were obtained for the genes of the clade B-E in *Drosophila* (Egli *et al.* 2003; Guirola *et al.* 2011). Those observations suggest that the MT content of insect genomes is mainly shaped by many lineage-specific duplication events.

**Figure 2.**
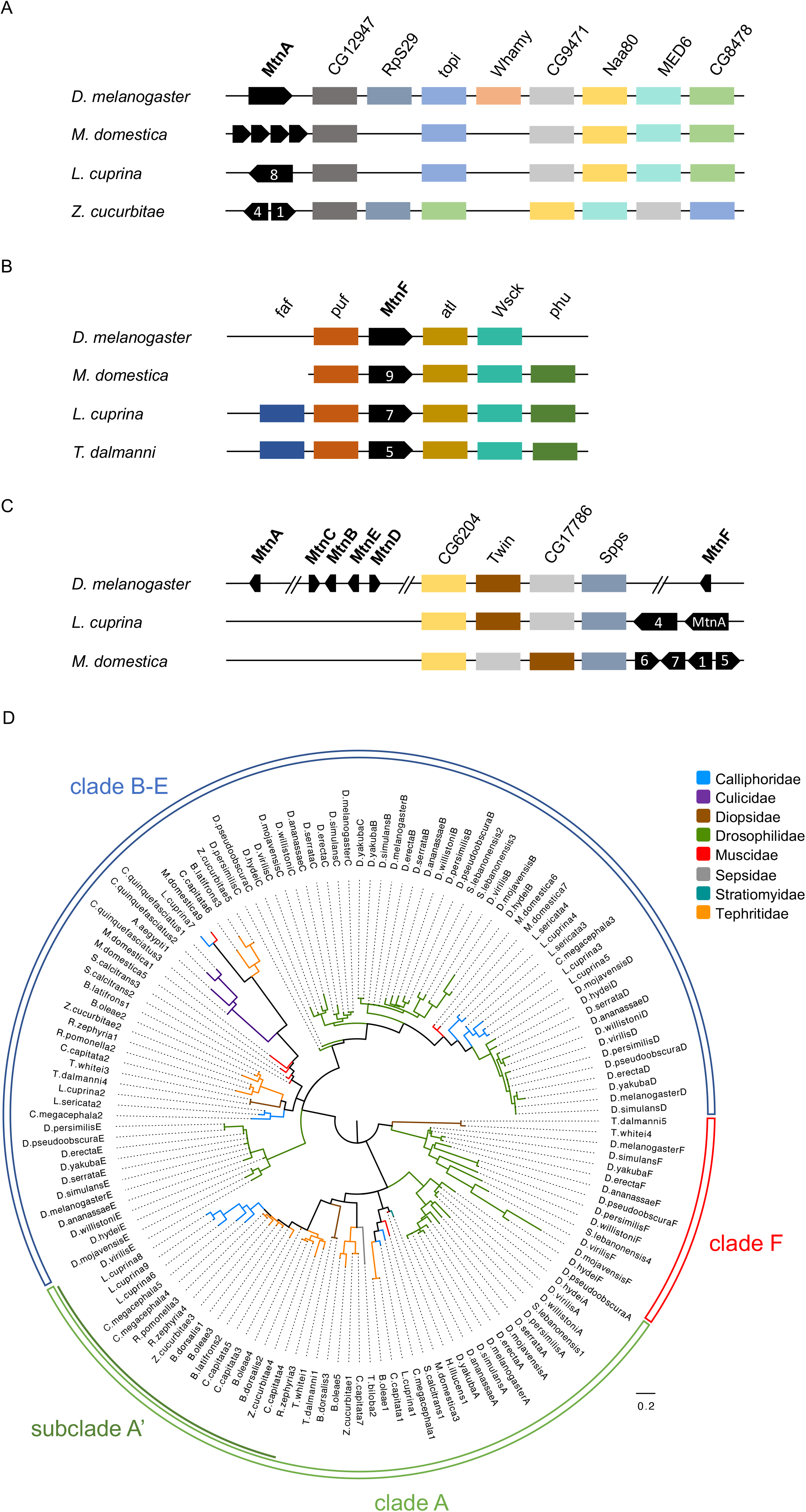
Synteny and phylogeny of MT genes in Diptera. (A) Deep conservation across the Diptera at the *MtnA* locus. (B) Deep conservation across the Diptera at the *MtnF* locus. (C) Conserved chromosomal clustering of the *MtnA*, *MtnB-E*, and *MtnF* loci. (D) The phylogram of the 141-taxon analyses was obtained through PhyML maximum-likelihood reconstruction. Analyses were conducted using the LG+Γ model. Support values obtained after 1,000 bootstrap replicates are shown in online supplementary information. Scale bar indicates number of changes per site.

Among the insects we investigated, lepidopteran insects show a high number of duplicated MTs with an average of 2.7 copies in tandem per species. The possibility of an ancient whole-genome duplication in lepidopteran insects has been firmly ruled out (Nakatani and McLysaght 2019). On the contrary, tandem duplications are associated with evolutionarily relevant traits in the butterflies and moths such as host plant detoxification (Fischer *et al.* 2008), gustatory and odorant receptors diversification (Briscoe *et al.* 2013; Pinharanda *et al.* 2017), vision (Smith and Briscoe 2015), wing color variation (Pinharanda *et al.* 2017; Westerman *et al.* 2018). Intensive tandem duplication and retention of MT genes we identified in the Lepidoptera reinforces the putative adaptive role of these proteins. Our findings raise the question about the specific high number of MT genes in lepidopteran insects. Caterpillars feed on plant leaves, but also on seeds and flowers of a large or restricted range of host species according to their generalist or specialist lifestyle. After metamorphosis, butterflies and moths feed primarily from flowers, and recent studies have shown that heavy metals can accumulate into floral organs and rewards (*i.e.*, nectar and pollen) (Milošević *et al.* 2014; Dhiman *et al.* 2017; Xun *et al.* 2017). In line with the increase in MT copies that correlates with augmented metal tolerance in *Drosophila* (Maroni *et al.* 1987; Otto *et al.* 1987; Meyer *et al.* 2006), having a richer palette of MT proteins might be an advantage for the fitness of Lepidoptera that occupy and feed on a wide variety of host species/tissues during their life-cycle.

### Evolution of the MT repertoire in the Diptera

As our large-scale analysis suggested substantial changes in the Diptera, we focused on the evolution of MTs in this particular insect order. We performed maximum-likelihood phylogenetic reconstruction using an alignment of 141 MT proteins restricted to the Diptera. The tree obtained clarifies the number of main MT clades and their relationships (**Figure 2D**). In this tree, MT sequences split into three main clades: clade A contains orthologues of the *D. melanogaster* gene *MtnA* (Lastowski-Perry *et al.* 1985), clade B-E contains homologues of the *D. melanogaster MtnB*-like cluster genes (Egli *et al.* 2003; Atanesyan *et al.* 2011) and clade F contains orthologues of the uncharacterized *D. melanogaster* ORF *CG43222*. Clade F, which represents a new clade of MT previously unidentified, is discussed in detail in the next section. Clade B-E contains sequences from the early-diverging Culicidae (mosquitoes), indicating that clade B-E can be traced back to the origin of Diptera. Clades A and F contain sequences that encompass several families of Diptera, suggesting that clades A and F could also be traced back to the origin of Diptera. According to the tree topology, it is reasonable to propose that the last common ancestor of extant Diptera possessed at least three MT genes.

We also investigated the conservation of synteny at the MT loci. Genomic data were available for eight sequences that groups into clade A in the species *D. melanogaster*, *M. domestica*, *Lucilia cuprina*, and *Zeugodacus cucurbitae*. All these MT sequences show a high level of synteny conservation with a recurrent handful of flanking genes (**Figures 2A-C**). Thus, these eight MT genes have the exact same relative genomic location in four different species. Our results indisputably confirm that sequences clustered in clade A correspond to orthologues of *MtnA*. We found such a pattern of conserved microsynteny for sequences within clade F (**Figure 2B**). The synteny of the genes of the *MtnB*-like cluster is less conserved at microscale, but well conserved at a larger genomic scale (**Figure 2C**). Our finding suggests that the numerous MT genes are clustered on the same part of the chromosome in at least *L. cuprina* (Calliphoridae) and *M. domestica* (Muscidae), as it is the case in Drosophilidae (Guirola *et al.* 2011). Our syntenic approach also confirms that the last common ancestor of extant Diptera possessed at least three MT genes.

### Identification of MtnF, a new MT member in the Diptera

Our phylogenetic analysis reveals a new clade of MT in the extant Diptera (**Figures 1A and 2D**). With regard to the MT repertoire already known in *D. melanogaster*, we subsequently named it *MtnF* in this species, in which it corresponds to the uncharacterized ORF *CG43222*. This finding is even more remarkable in the model species *D. melanogaster* as the MT family has been studied since 1980s. Following the cloning of its first MT (Lastowski-Perry *et al.* 1985), *D. melanogaster* genome had been reported to contain four MT genes (*MtnA*, *MtnB*, *MtnC* and *MtnD*) that are clustered in the right arm of the third chromosome (Egli *et al.* 2003). In 2011, the fifth MT gene *MtnE* was identified and shown to locate inside the MtnB-like cluster in *D. melanogaster* (Atanesyan *et al.* 2011).

The alignment of the N-terminal end of MtnF with its *Drosophila* homologs highlights the conservation between the different MT members (**Figure 3A**), suggesting that MtnF is a member of the MT family. To better characterize this new MT member, we predicted the 3D structure of the protein MtnF in *D. melanogaster*. The best model (C-score = −0.63) predicts two possible β-sheet secondary structures (**Figure 3B**) and fits well (TM-score = 0.538) to the crystal structure of the rat Mt2 (purple structure, **Figures 3C-D**), supporting the hypothesis that MtnF might function as a MT. Another piece of evidence supporting *MtnF* as a MT-encoding gene is the presence of two putative MTF-1 binding sites in its upstream region (**Figure S1**). We found one core consensus sequence of metal response element TGCRCNCG (Günther *et al.* 2012; Sims *et al.* 2012) in the non-coding 5’ region of the *MtnF* locus, and one in the coding region of the 5’ flanking gene *puffyeye*.

**Figure 3.**
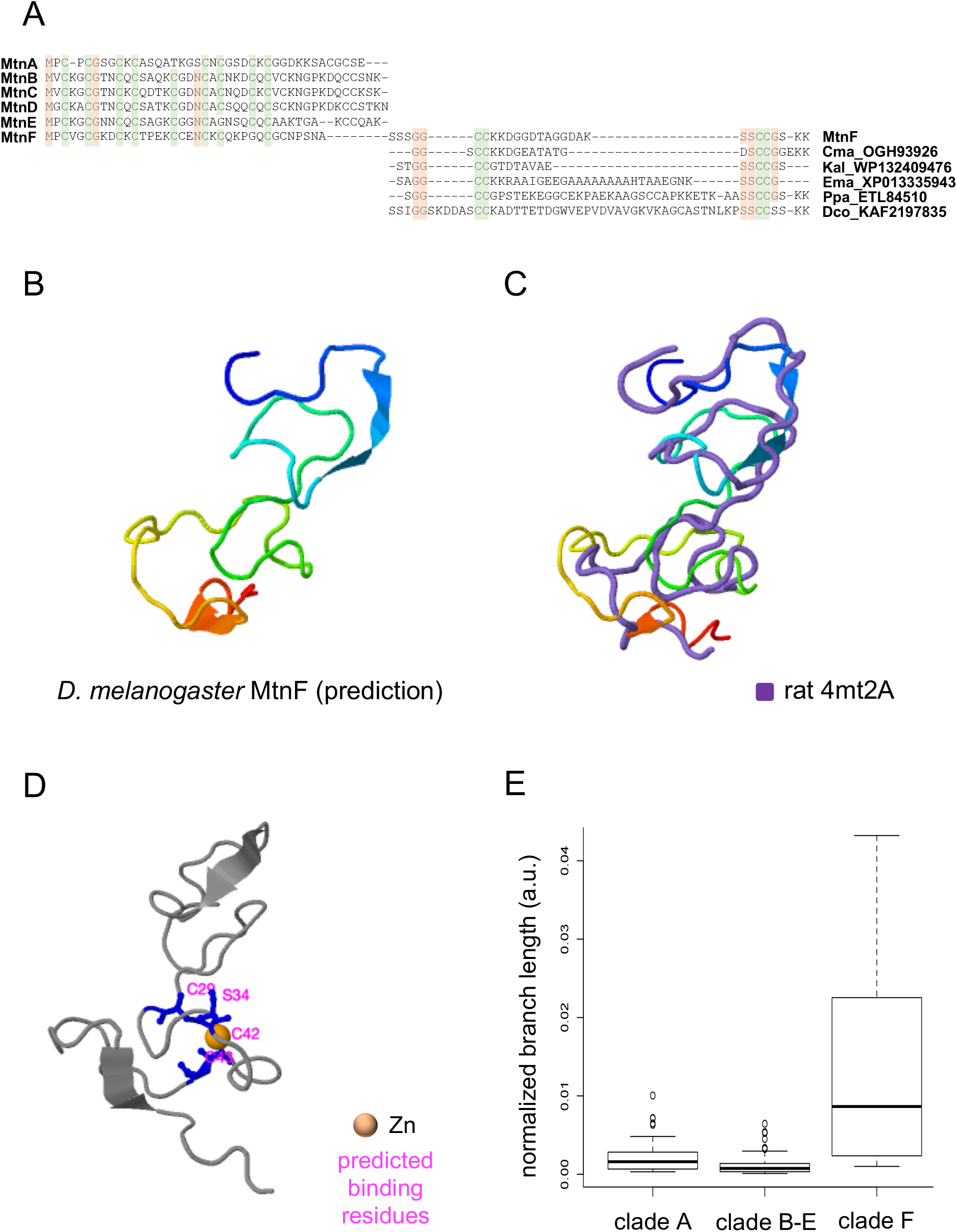
Evolution and structure of the newly identified MtnF of *D. melanogaster*. (A) Alignment of the six MT proteins of *D. melanogaster*. MtnF has an extra motif in C-terminus that can be aligned with non-insect proteins. (B) Predicted 3D conformation of the protein MtnF of *D. melanogaster* (RasMol colors). (C) Alignment of the predicted 3D structure of *D. melanogaster* MtnF with the one of the rat MT 4mt2A obtained by crystallography. (D) Position of the atom of Zn predicted as ligand and predicted associated binding residues. (E) Comparison of branch lengths between clade A, clade B-E, and clade F. Cma: *Candidatus magasanikbacteria*, Kba: *Kribbella albertanoniae*, Ema: *Eimeria maxima*, Ppa: *Phytophthora parasitica*, Dco: *Delitschia confertaspora*.

#### Might MtnF be a Zn-thionein in the Diptera?

As previously mentioned, a striking feature is the absence of Zn-thioneins in insects whereas they are found in all metazoans (Atrian 2009). The identification of a sixth MT member in *D. melanogaster* naturally raises the question of whether MtnF could act as a Zn-thionein. We found that *MtnF* orthologs evolve faster than *MtnA* and *MtnB-E* orthologs. On average, *MtnF* orthologs have longer branch lengths than *MtnA* (*t*-test: df = 23, P = 0.0003) and *MtnB-E* (*t*-test: df = 23, P = 0.0001) orthologs (**Figure 3E**). This higher divergence might explain why previous searches failed to identify *CG43222* as a MT-coding gene.

With an average length of 62 amino acids the proteins of the clade F are the longest MTs in insects (**Figure 4A**). The alignment of the MtnF protein sequence with the other five MTs in *D. melanogaster* shows the overlength is due to extra amino acid residues in C-terminus (**Figure 3A**). This observation raises the question of the origin of these residues. BLASTP and PSI-BLAST searches identified similar motifs in bacteria (*Candidatus magasanikbacteria*, *Kribbella albertanoniae*), apicomplexan protozoans (*Eimeria maxima*), fungus-like oomycota (*Phytophthora parasitica*), and fungi (*Delitschia confertaspora*). Interestingly, the proteins Ema_XP013335943 and Ppa_ETL84510 are a putative copper transporter and a putative heavy-metal-translocating P-type ATPase, respectively. However, the conservation of the N-terminus across *Drosophila* MTs does not really support an acquisition of the C-term peptide by horizontal transfer.

**Figure 4.**
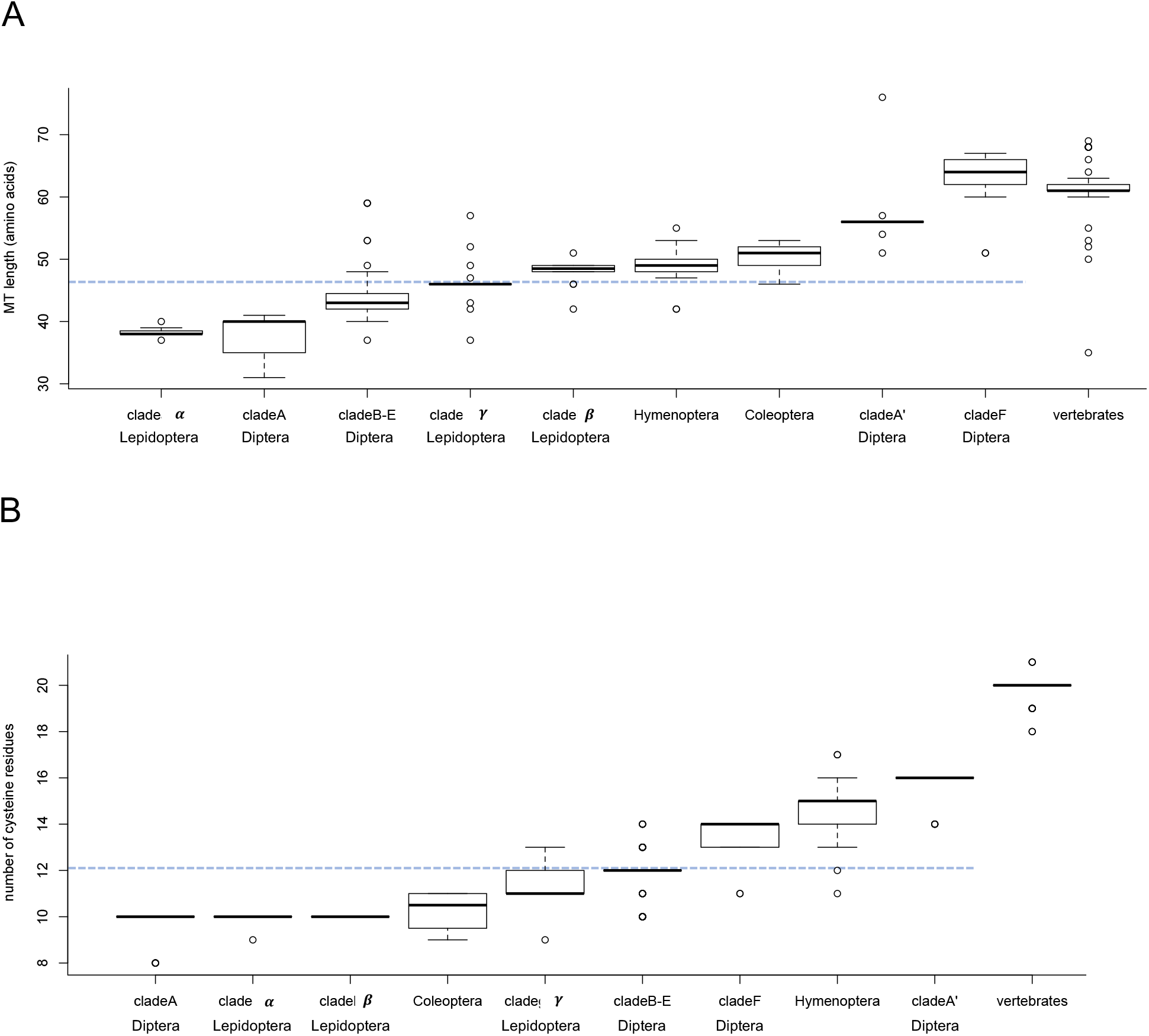
Comparative characterization of two MT features. (A) Length of MT proteins in insects and vertebrates, (B) Number of cysteine residues per MT in insects and vertebrates. The blue dotted line indicates the mean of the plotted variable.

The augmented length of the clade F proteins goes along with a richer content in cysteine. We counted the number of cysteines per MT protein for each clade (**Figure 4B**). On average, proteins of the MtnF clade contain 13.4 cysteines, which is statistically higher than the average number of 12 cysteines found in insects (*t*-test: df = 13, P < 1.4e-04). We also unraveled a bimodal distribution of the number of cysteines within the clade A. Whereas most of the clade A proteins contain an average number of 10 cysteines (**Figure 4B**), a subgroup of sequences, subclade A’ contains an average number of 16 cysteines (**Figure 4B**) that is statistically higher than the cysteine average content in the clade A (t-test: df = 17, P < 1.0e-05) and in insects (t-test: df = 17, P < 1.0e-05). The embedded position of the clade A’ within the clade A suggests that the average gain of 6 cysteines results from a unique event during the evolution of the clade A. Similarly, we observed a cysteine enrichment in the hymenopteran MTs (*t*-test: df = 33, P < 1.0e-05) compared to the average cysteine content in insects. Our study suggests that the enrichment in cysteine is a derived MT feature that appeared several times independently during the course of evolution.

What could be the biological function(s) of MtnF? Determining the specificity of metal binding properties of MTs has been a prevalent question for the last years. Because of their longer size and higher cysteine content, the MTs of the clade F are reminiscent of the vertebrate MTs (**Figures 4A and 4B**). Vertebrate MTs contain two metal-thiolate clusters named β-domain with three binding sites for divalent ions involving 9 cysteinyl sulphurs, and α-domain capable of binding four divalent metal ions involving 11 cysteinyl sulphurs (Robbins *et al.* 1991; Rigby Duncan and Stillman 2007), and are able to bind divalent Zn(II) contrary to insect MTs. According to its structural and evolutionary peculiarities, we propose that MtnF might be a putative Zn-thionein in the Diptera. According to the database FlyAtlas2, the highest expression of *MtnF* has been reported in the *Drosophila* testis (Leader *et al.* 2018). The protein MtnF is also more expressed in malignant brain tumors in females than in males (Molnar *et al.* 2019), reinforcing the idea that this gene can have a sex-biased expression. Another common feature of *MtnF* is its expression in nervous system cells like sensilla hosting olfactory receptors neurons (Mohapatra and Menuz 2019), brain cells and neurons in *D. melanogaster* (Crocker *et al.* 2016), especially cholinergic neurons (Avalos *et al.* 2019). Last, a recent comparative study shows that heat stress induces transcription of *MtnF* more in the sensitive *D. hydei* than in the tolerant *D. buzzatii* (Rane *et al.* 2019).

#### Additional peculiarities of dipteran MTs

In rare cases, histidine residues have been found to coordinate divalent metals instead of cysteines. Histidine coordination is carried out by SmtA in the cyanobacterium *Synechococcus* PCC7942 (Daniels *et al.* 1998) and by Ec in the wheat (Leszczyszyn *et al.* 2007). Moreover, histidine residues have been found in a variety of MTs (Blindauer and Leszczyszyn 2010). These results prompted us to investigate the histidine content of insect MTs. We found 31 sequences with one histidine and 6 sequences with two histidines, but such a limited number of histidines might not be relevant for metal coordination. More interestingly, we found one sequence with three histidines (*M.domestica1*) and one sequence with five histidines (*M.domestica8*) in the housefly *M. domestica*. The larva of the housefly is known for expressing MTs that have an antibacterial activity (Jin *et al.* 2005). It would be interesting to test whether the rich histidine content contribute to the antibacterial activity.

## CONCLUSION

In this paper, we identified more than 300 new insect MT sequences and studied their evolutionary relationship using a combination of phylogenetics and synteny analysis. This combined approach allowed us to leverage the conserved microsynteny in insects to reconstruct the evolution of sequences with limited informative sites. Our data suggest that the presence of a single MT in the last common ancestor of extant insects (**Figure 5**). The MT content was subsequently shaped by lineage-specific tandem duplications. In particular, a gene duplication event predated the divergence between Diptera and Lepidoptera (**Figure 5**). We suggest that the dynamic changes in the MT content across insects may reflect differences in environment, diet, and past contact with heavy metals. We also expect that the newly identified MT sequences will allow to develop future biomarkers of heavy metal contamination. More importantly, our combined phylogenetics and synteny analysis can allow us to identify other previously unidentified genes and other low complexity genomic features.

**Figure 5.**
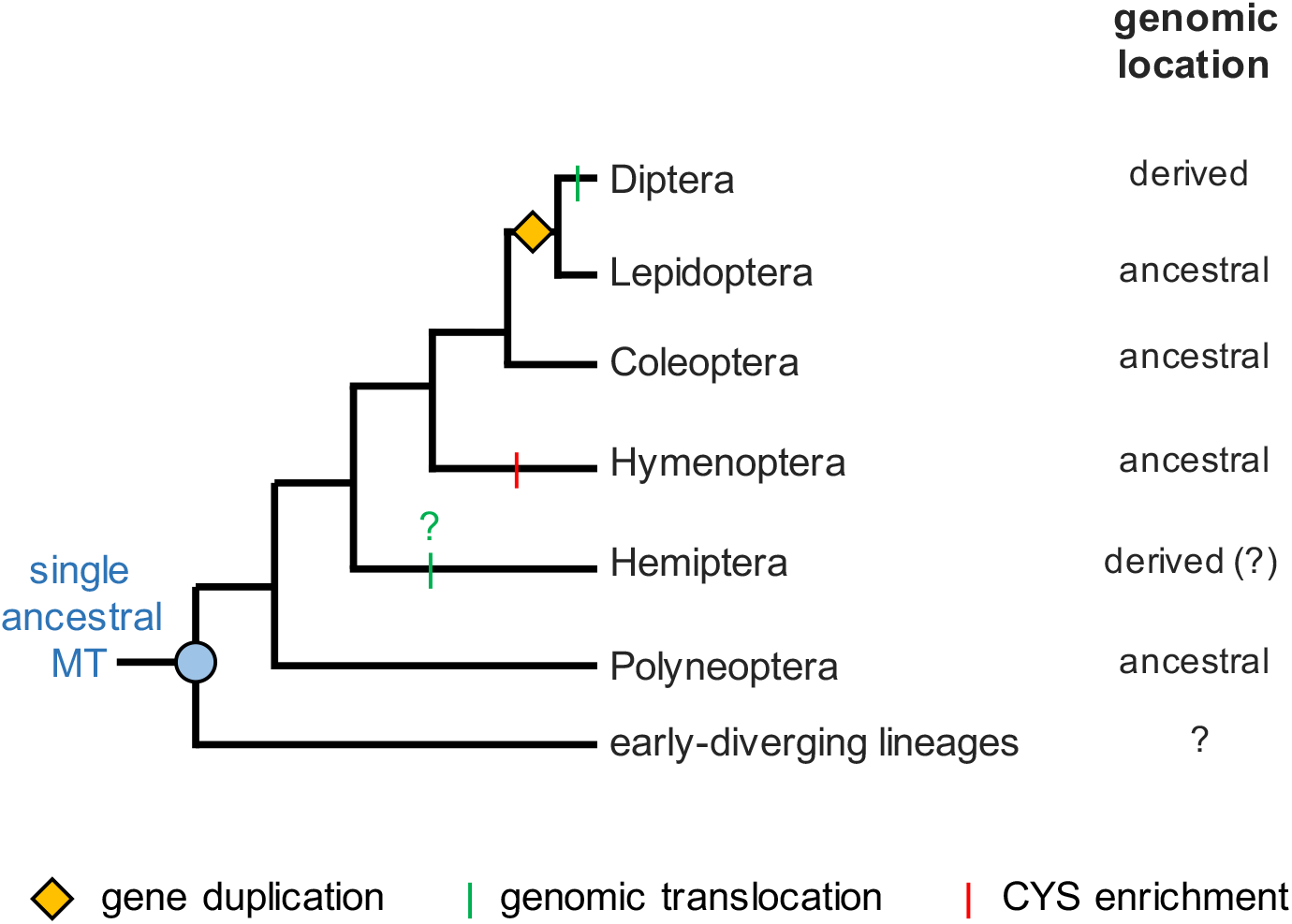
Summary of main molecular events during insect evolution. Gene duplications events are depicted by a yellow diamond, genomic translocations by a vertical green bar, and cysteine-enrichment by a vertical red line.

## MATERIALS AND METHODS

### Data collection

MT sequences were identified in 116 insect transcriptomes by TBLASTN using 20 published MT sequences as queries (Table S1). The transcriptomes were retrieved from the InsectBase website (http://www.insect-genome.com/) (Yin *et al.* 2015) and used to build a local database with BioEdit 5.0.6 (Hall 1999). The putative identified MTs were added to the set of probes, and TBLASTN searches were iteratively repeated until no new MTs were found. Subsequently, we used all these newly identified MTs as probes to search for additional MT sequences in NCBI NR (protein) and NT (nucleotide) databases using BLASTP and TBLASTN, respectively. In order to remove false positive MTs, we predicted open reading frames (ORFs) of all putative MT nucleotide sequences using the EMBOSS program getorf (http://www.bioinformatics.nl/cgi-bin/emboss/getorf). Putative MT sequences for which we failed to identify ORFs were discarded. Finally, redundant copies retrieved from InsectBase, NCBI NR and NT databases were manually removed from our dataset. Species full names are the following ones: *Acromyrmex echinatior*, *Aedes aegypti*, *Aethina tumida*, *Anthonomus grandis*, *Apis florea*, *A. mellifera*, *Aretaon asperrimus*, *Asbolus verrucosus*, *Bactrocera dorsalis*, *B. latifrons*, *B. oleae*, *Bicyclus anynana*, *Bombus impatiens*, *B. terrestris*, *Bombyx mandarina*, *B. mori*, *Camponotus floridanus*, *Cephus cinctus*, *Ceratitis capitata*, *Ceratosolen solmsi*, *Chilo suppressalis*, *Chrysomya megacephala*, *Chyphotes mellipes*, *Cnaphalocrocis medinalis*, *Copidosoma floridanum*, *Cotesia vestalis*, *Cryptotermes secundus*, *Culex quinquefasciatus*, *Cyphomyrmex costatus*, *Danaus plexippus*, *Dendroctonus frontalis*, *Diachasma alloeum*, *Drosophila ananassae*, *D. erecta*, *D. hydei*, *D. melanogaster*, *D. mojavensis*, *D. persimilis*, *D. pseudoobscura*, *D. serrata*, *D. simulans*, *D. virilis*, *D. willistoni*, *yakuba*, *Dufourea novaeangliae*, *Eufriesea mexicana*, *Extatosoma tiaratum*, *Fopius arisanus*, *Galleria mellonella*, *Gryllus bimaculatus*, *Heliconius melpomene*, *Helicoverpa armigera*, *Hermetia illucens*, *Hyposmocoma kahamanoa*, *Laphygma exigua*, *Leptopilina boulardi*, *Leptopilina heterotoma*, *Lucilia cuprina*, *Lucilia sericata*, *Manduca sexta*, *Medauroidea extradentata*, *Microplitis demolitor*, *Mischocyttarus flavitarsis*, *Musca domestica*, *Ooceraea biroi*, *Onthophagus nigriventris*, *Orussus abietinus*, *Osmia cornuta*, *Ostrinia furnacalis*, *O. scapulalis*, *Oxya chinensis*, *Papilio machaon*, *Papilio polytes*, *Papilio xuthus*, *Pieris rapae*, *Pogonomyrmex barbatus*, *Polistes canadensis*, *Polistes metricus*, *Rhagoletis pomonella*, *Rhagoletis zephyria*, *Scaptodrosophila lebanonensis*, *Sceliphron caementarium*, *Sipyloidea sipylus*, *Sphaerophtalma orestes*, *Spodoptera exigua*, *Spodoptera litura*, *Stomoxys calcitrans*, *Teleopsis dalmanni*, *Teleopsis whitei*, *Themira biloba*, *Trachymyrmex cornetzi*, *Trachymyrmex septentrionalis*, *Trialeurodes vaporariorum*, *Tribolium castaneum*, *Trichogramma pretiosum*, *Trichoplusia ni*, *Tryporyza incertulas*, *Trypoxylus dichotomus*, *Vanessa tameamea*, *Zeugodacus cucurbitae*, *Zootermopsis nevadensis*, *Zygaena filipendulae*. Datasets, sequence alignments, and tree files are downloadable from Dryad (https://datadryad.org/stash/share/0WQ8Btoh6abEUsEIBUV4RMkL52XzaCBp0wG0Av8mEWY).

### Phylogenetic analysis

Amino acid sequences were aligned with MUSCLE (Edgar 2004), manually adjusted, and selected blocks were used for phylogenetic reconstruction. Maximum-likelihood searches were performed using PhyML 3.0 (Guindon *et al.* 2010) under the LG substitution matrix with final likelihood evaluation using a gamma distribution. 1,000 bootstrap replicates were conducted for support estimation.

### Synteny analysis

Genomic sequences were retrieved from NCBI (http://www.ncbi.nlm.nih.gov), InsectBase (Yin *et al.* 2015), and 5,000 Insect Genome Project (i5k) (Poelchau *et al.* 2018) databases by BLASTN using the corresponding coding sequences as queries. Gene and order content of the genomic scaffolds or contigs were assessed by BLASTX against the annotated proteins of *D. melanogaster* (release 6.31)(Thurmond *et al.* 2018).

### Statistical analysis

Branch lengths (BL) were obtained as outputs of PhyML software (Guindon *et al.* 2010). To consider differences in number of sequences per clade, we calculated the normalized BL, that is the value of BL/number of MT sequences per clade. We compared the BL means between clade A, clade B-E, and clade F using a t-test as the BL followed a normal distribution. Statistical tests and graphics were performed using R statistics package version 3.5.0 (the R Project for Statistical Computing, www.r-project.org, last accessed 2019 December 17).

### Prediction of protein structure

The 3D structure of the newly identified metallothionein *MtnF* in *D. melanogaster* was predicted using the I-TASSER server (Yang and Zhang 2015). I-TASSER simulations generate a large ensemble of structural conformations, called decoys. To select the final models, I-TASSER uses the SPICKER program to cluster all the decoys based on the pairwise structure similarity and reports up to five models which corresponds to the five largest structure clusters. The confidence of each model is quantitatively measured by C-score that is calculated based on the significance of threading template alignments and the convergence parameters of the structure assembly simulations. C-score is typically in the range of [−5, 2], where a C-score of a higher value signifies a model with a higher confidence. The crystal structure of the rat Mt2 (Braun *et al.* 1992) (PDB code 4mt2) was used to model the MtnF protein. Ligand binding sites were predicted using the COFACTOR software (Zhang *et al.* 2017).

## ACKNOWLEDGEMENT

This work is supported by USDA NIFA via Michigan State University AgBioresearch (Umbrella project MICL02522 to HC) and by the National Natural Science Foundation of China (No. 31760637, 31640064 to HYW). ML is supported by the China Scholarship Council (CSC) (File No. 201708360095).

## AUTHOR CONTRIBUTIONS

**Conceptualization:** ML, HC

**Acquisition of Data:** ML, HSC

**Analysis of Data:** CF, ML, HC

**Funding acquisition:** HC and HYW

**Project administration**: HC

**Wrote the initial manuscript**: CF, HC

**Contributed to subsequent drafts of the manuscript**: ML, CF, HSC, HYW, HC.

## Supplementary figure and table legends

**Table S1.** Summary of identified MTs prior to our study.

| Class | Order | Family | Species | Name | Accession | Reference |
| --- | --- | --- | --- | --- | --- | --- |
| Insecta | Diptera | Chironomidae | *Chironomus riparius* | *C.riparius1* | HQ260607 | Park and Kwak 2012 |
|  | Diptera | Culicidae | *Anopheles gambiae* | *A.gambiae1* | AY994093 | Arcà *et al.* 2005 |
|  | Diptera | Culicidae | *Anopheles gambiae* | *A.gambiae2* | AY994094 | Arcà *et al.* 2005 |
|  | Diptera | Drosophilidae | *Drosophila melanogaster* | *D.melanogasterA* | NM079575 | Lastowski-Perry *et al.* 1985 |
|  | Diptera | Drosophilidae | *Drosophila melanogaster* | *D.melanogasterB* | NM079689 | Silar *et al.* 1990 |
|  | Diptera | Drosophilidae | *Drosophila melanogaster* | *D.melanogasterC* | NM142625 | Egli *et al.* 2003 |
|  | Diptera | Drosophilidae | *Drosophila melanogaster* | *D.melanogasterD* | NM176518 | Egli *et al.* 2003 |
|  | Diptera | Drosophilidae | *Drosophila melanogaster* | *D.melanogasterE* | NM001202325 | Atanesyan *et al.* 2011 |
|  | Diptera | Muscidae | *Musca domestica* | *M.domestica2* | JF919737 | Tang *et al.* 2011 |
|  | Diptera | Tabanidae | *Tabanus yao* | *T.yao1* | EU147261 | Xu *et al.* 2008 |
|  | Diptera | Tabanidae | *Tabanus yao* | *T.yao2* | EU147262 | Xu *et al.* 2008 |
|  | Hymenoptera | Apidae | *Apis florea* | *A.florea1* | XP003689576 | Purać *et al.* 2019 |
|  | Hymenoptera | Apidae | *Apis mellifera* | *A.mellifera1* | XM001120071 | Purać *et al.* 2019 |
|  | Hymenoptera | Apidae | *Bombus impatiens* | *B.impatiens1* | XR001102067 | Purać *et al.* 2019 |
|  | Hymenoptera | Apidae | *Bombus impatiens* | *B.impatiens2* | XR002947883 | Purać *et al.* 2019 |
|  | Hymenoptera | Apidae | *Bombus terrestris* | *B.terrestris1* | XR001099390 | Purać *et al.* 2019 |
|  | Hymenoptera | Apidae | *Bombus terrestris* | *B.terrestris2* | XM020866906 | Purać *et al.* 2019 |
|  | Orthoptera | Acridoidea | *Oxya chinensis* | *O.chinensis1* | KJ153014 | Liu *et al.* 2014 |
|  | Orthoptera | Acridoidea | *Oxya chinensis* | *O.chinensis2* | KJ153015 | Liu *et al.* 2014 |
| Entognatha | Collembola | Entomobryidae | *Orchesella cincta* | *O.cincta1* | AAD10193 | Hensbergen *et al.* 1999 |

**Figure S1.**
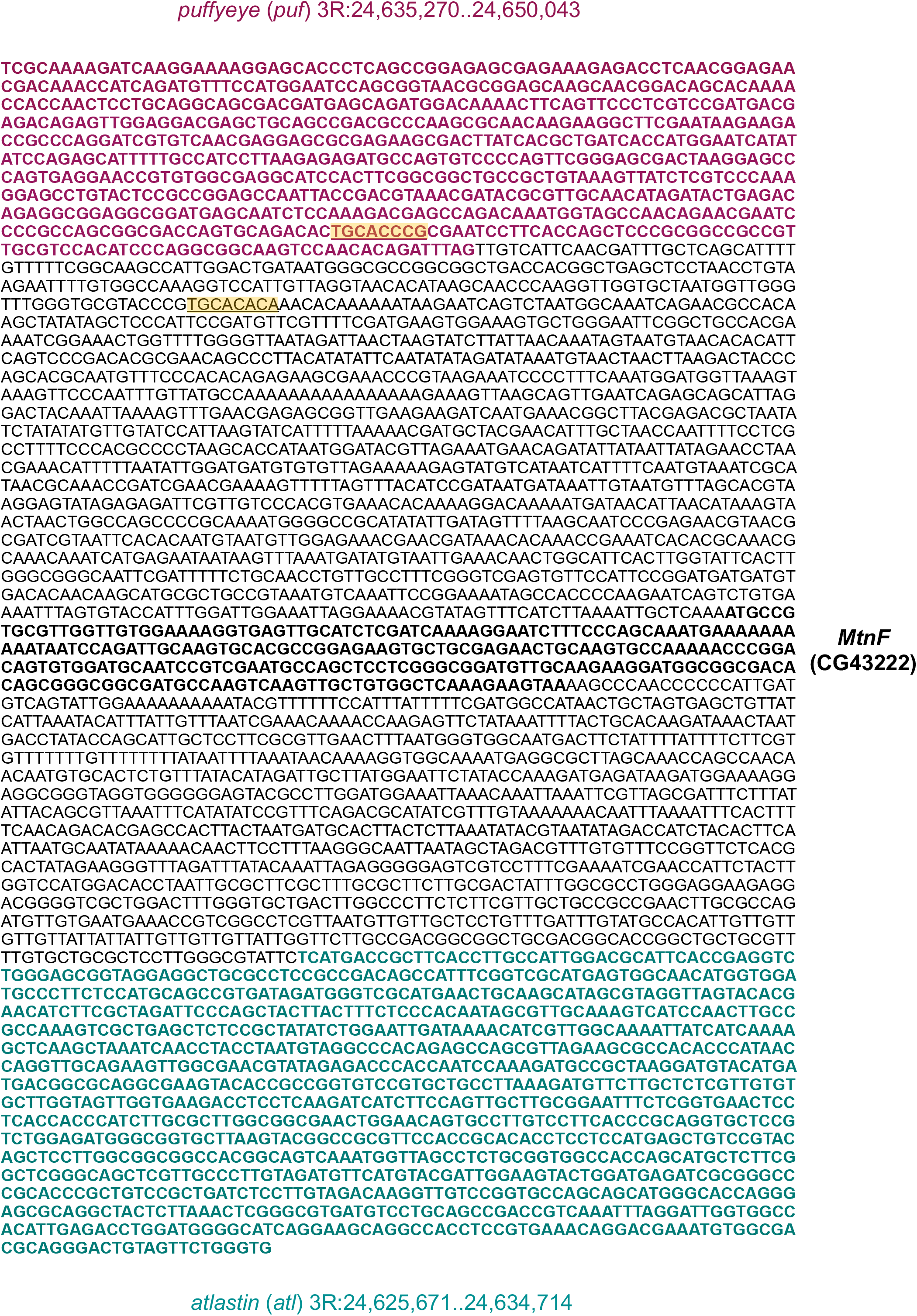
Putative MTF-1 binding sites upstream *MtnF* in *D. melanogaster*. The MTF-1 binding sites are highlighted in yellow.

## REFERENCES

Adams, M. D., S. E. Celniker, R. A. Holt, C. A. Evans, J. D. Gocayne, P. G. Amanatides, S. E. Scherer, P. W. Li, R. A. Hoskins and R. F. Galle, 2000 The genome sequence of Drosophila melanogaster. Science 287:2185–2195.

Adler, P. H., O. Yadamsuren and W. S. Procunier, 2016 Chromosomal translocations in black flies (Diptera: Simuliidae)—facilitators of adaptive radiation? PloS one 11.

Artemov, G. N., A. N. Peery, X. Jiang, Z. Tu, V. N. Stegniy, M. V. Sharakhova and I. V. Sharakhov, 2017 The physical genome mapping of Anopheles albimanus corrected scaffold misassemblies and identified interarm rearrangements in genus Anopheles. G3: Genes, Genomes, Genetics 7:155–164.

Atanesyan, L., V. Günther, S. E. Celniker, O. Georgiev and W. Schaffner, 2011 Characterization of MtnE, the fifth metallothionein member in Drosophila. JBIC Journal of Biological Inorganic Chemistry 16:1047.

Atrian, S., 2009 6: Metallothioneins in Diptera.

Avalos, C. B., G. L. Maier, R. Bruggmann and S. G. Sprecher, 2019 Single cell transcriptome atlas of the Drosophila larval brain. ELife 8.

Binz, P.-A., and J. H. Kägi, 1999 Metallothionein: molecular evolution and classification, pp. 7–13 in Metallothionein Iv. Springer.

Blindauer, C. A., and O. I. Leszczyszyn, 2010 Metallothioneins: unparalleled diversity in structures and functions for metal ion homeostasis and more. Natural product reports 27:720–741.

Braun, W., M. Vasak, A. Robbins, C. Stout, G. Wagner, J. Kägi and K. Wüthrich, 1992 Comparison of the NMR solution structure and the x-ray crystal structure of rat metallothionein-2. Proceedings of the National Academy of Sciences 89:10124–10128.

Briscoe, A. D., A. Macias-Munoz, K. M. Kozak, J. R. Walters, F. Yuan, G. A. Jamie, S. H. Martin, K. K. Dasmahapatra, L. C. Ferguson and J. Mallet, 2013 Female behaviour drives expression and evolution of gustatory receptors in butterflies. PLoS genetics 9.

Calvete, O., J. González, E. Betrán and A. Ruiz, 2012 Segmental duplication, microinversion, and gene loss associated with a complex inversion breakpoint region in Drosophila. Molecular biology and evolution 29:1875–1889.

Capdevila, M., and S. Atrian, 2011 Metallothionein protein evolution: a miniassay. JBIC Journal of Biological Inorganic Chemistry 16:977–989.

Cobbett, C., and P. Goldsbrough, 2002 Phytochelatins and metallothioneins: roles in heavy metal detoxification and homeostasis. Annual review of plant biology 53:159–182.

Crocker, A., X.-J. Guan, C. T. Murphy and M. Murthy, 2016 Cell-type-specific transcriptome analysis in the Drosophila mushroom body reveals memory-related changes in gene expression. Cell reports 15:1580–1596.

d’Alencon, E., H. Sezutsu, F. Legeai, E. Permal, S. Bernard-Samain, S. Gimenez, C. Gagneur, F. Cousserans, M. Shimomura and A. Brun-Barale, 2010 Extensive synteny conservation of holocentric chromosomes in Lepidoptera despite high rates of local genome rearrangements. Proceedings of the National Academy of Sciences 107:7680–7685.

Daniels, M. J., J. S. Turner-Cavet, R. Selkirk, H. Sun, J. A. Parkinson, P. J. Sadler and N. J. Robinson, 1998 Coordination of Zn2+ (and Cd2+) by Prokaryotic Metallothionein INVOLVEMENT OF HIS-IMIDAZOLE. Journal of Biological Chemistry 273:22957–22961.

Dhiman, S. S., X. Zhao, J. Li, D. Kim, V. C. Kalia, I.-W. Kim, J. Y. Kim and J.-K. Lee, 2017 Metal accumulation by sunflower (Helianthus annuus L.) and the efficacy of its biomass in enzymatic saccharification. PloS one 12.

Edgar, R. C., 2004 MUSCLE: a multiple sequence alignment method with reduced time and space complexity. BMC Bioinformatics 5:113.

Egli, D., J. Domènech, A. Selvaraj, K. Balamurugan, H. Hua, M. Capdevila, O. Georgiev, W. Schaffner and S. Atrian, 2006 The four members of the Drosophila metallothionein family exhibit distinct yet overlapping roles in heavy metal homeostasis and detoxification. Genes to Cells 11:647–658.

Egli, D., A. Selvaraj, H. Yepiskoposyan, B. Zhang, E. Hafen, O. Georgiev and W. Schaffner, 2003 Knockout of ‘metal‐responsive transcription factor’MTF‐1 in Drosophila by homologous recombination reveals its central role in heavy metal homeostasis. The EMBO journal 22:100–108.

Finet, C., A. Decaras, D. Armisen and A. Khila, 2018 The achaete–scute complex contains a single gene that controls bristle development in the semi-aquatic bugs. Proceedings of the Royal Society B 285:20182387.

Finet, C., K. Slavik, J. Pu, S. B. Carroll and H. Chung, 2019 Birth-and-death evolution of the fatty acyl-CoA reductase (FAR) gene family and diversification of cuticular hydrocarbon synthesis in *Drosophila*. Genome Biol Evol 11:1541–1551.

Fischer, H. M., C. W. Wheat, D. G. Heckel and H. Vogel, 2008 Evolutionary origins of a novel host plant detoxification gene in butterflies. Molecular Biology and Evolution 25:809–820.

Guindon, S., J.-F. Dufayard, V. Lefort, M. Anisimova, W. Hordijk and O. Gascuel, 2010 New algorithms and methods to estimate maximum-likelihood phylogenies: assessing the performance of PhyML 3.0. Systematic biology 59:307–321.

Guirola, M., Y. Naranjo, M. Capdevila and S. Atrian, 2011 Comparative genomics analysis of metallothioneins in twelve Drosophila species. Journal of inorganic biochemistry 105:1050–1059.

Günther, V., U. Lindert and W. Schaffner, 2012 The taste of heavy metals: gene regulation by MTF-1. Biochimica et Biophysica Acta (BBA)-Molecular Cell Research 1823:1416–1425.

Hall, T. A., 1999 BioEdit: a user-friendly biological sequence alignment editor and analysis program for Windows 95/98/NT, pp. 95–98 in Nucleic acids symposium series. [London]: Information Retrieval Ltd., c1979-c2000.

Hamer, D. H., 1986 Metallothionein. Annual review of biochemistry 55:913–951.

Janssens, T. K., D. Roelofs and N. M. Van Straalen, 2009 Molecular mechanisms of heavy metal tolerance and evolution in invertebrates. Insect Science 16:3–18.

Jin, H., Q. Hui, J. Jun, A. C. Ju, L. Sen, D. Qian and Q. Lin, 2005 Preliminary studies on the zinc-induced metallothionein protein with antibacterial activity in housefly larvae, Musca domestica. Acta Biologica Hungarica 56:283–295.

Klaassen, C. D., J. Liu and S. Choudhuri, 1999 Metallothionein: an intracellular protein to protect against cadmium toxicity. Annual review of pharmacology and toxicology 39:267–294.

Lastowski-Perry, D., E. Otto and G. Maroni, 1985 Nucleotide sequence and expression of a Drosophila metallothionein. Journal of Biological Chemistry 260:1527–1530.

Leader, D. P., S. A. Krause, A. Pandit, S. A. Davies and J. A. T. Dow, 2018 FlyAtlas 2: a new version of the Drosophila melanogaster expression atlas with RNA-Seq, miRNA-Seq and sex-specific data. Nucleic acids research 46:D809–D815.

Leszczyszyn, O. I., R. Schmid and C. A. Blindauer, 2007 Toward a property/function relationship for metallothioneins: Histidine coordination and unusual cluster composition in a zinc‐metallothionein from plants. Proteins: Structure, Function, and Bioinformatics 68:922–935.

Liu, Y., H. Wu, L. Kou, X. Liu, J. Zhang, Y. Guo and E. Ma, 2014 Two metallothionein genes in Oxya chinensis: molecular characteristics, expression patterns and roles in heavy metal stress. PloS one 9:e112759.

Mandrioli, M., G. Melchiori, M. Panini, O. Chiesa, R. Giordano, E. Mazzoni and G. C. Manicardi, 2019 Analysis of the extent of synteny and conservation in the gene order in aphids: A first glimpse from the Aphis glycines genome. Insect biochemistry and molecular biology 113:103228.

Maroni, G., J. Wise, J. Young and E. Otto, 1987 Metallothionein gene duplications and metal tolerance in natural populations of Drosophila melanogaster. Genetics 117:739–744.

Merritt, T. J., and A. J. Bewick, 2017 Genetic diversity in insect metal tolerance. Frontiers in genetics 8:172.

Meyer, J. L., M. A. Hoy and A. Jeyaprakash, 2006 Insertion of a yeast metallothionein gene into the model insect Drosophila melanogaster (Diptera: Drosophilidae) to assess the potential for its use in genetic improvement programs with natural enemies. Biological Control 36:129–138.

Milošević, T., M. Đurić and N. Milošević, 2014 Accumulation of Heavy Metals in Flowers of Fruit Species. Water, Air, & Soil Pollution 225:2019.

Misof, B., S. Liu, K. Meusemann, R. S. Peters, A. Donath, C. Mayer, P. B. Frandsen, J. Ware, T. Flouri and R. G. Beutel, 2014 Phylogenomics resolves the timing and pattern of insect evolution. Science 346:763–767.

Mohapatra, P., and K. Menuz, 2019 Molecular profiling of the Drosophila antenna reveals conserved genes underlying olfaction in insects. G3: Genes, Genomes, Genetics 9:3753–3771.

Mokdad, R., A. Debec and M. Wegnez, 1987 Metallothionein genes in Drosophila melanogaster constitute a dual system. Proceedings of the National Academy of Sciences 84:2658–2662.

Molnar, C., J. P. Heinen, J. Reina, S. Llamazares, E. Palumbo, A. Breschi, M. Gay, L. Villarreal, M. Vilaseca and G. Pollarolo, 2019 The histone code reader PHD finger protein 7 controls sex-linked disparities in gene expression and malignancy in Drosophila. Science Advances 5:eaaw7965.

Nakatani, Y., and A. McLysaght, 2019 Macrosynteny analysis shows the absence of ancient whole-genome duplication in lepidopteran insects. Proceedings of the National Academy of Sciences 116:1816–1818.

Navarro, J. A., and S. Schneuwly, 2017 Copper and zinc homeostasis: lessons from Drosophila melanogaster. Frontiers in genetics 8:223.

Otto, E., J. Allen, J. Young, R. Palmiter and G. Maroni, 1987 A DNA segment controlling metal-regulated expression of the Drosophila melanogaster metallothionein gene Mtn. Molecular and cellular biology 7:1710–1715.

Palacios, Ò., S. Atrian and M. Capdevila, 2011 Zn-and Cu-thioneins: a functional classification for metallothioneins? JBIC Journal of Biological Inorganic Chemistry 16:991.

Pérez-Rafael, S., A. Kurz, M. Guirola, M. Capdevila, Ò. Palacios and S. Atrian, 2012 Is MtnE, the fifth Drosophila metallothionein, functionally distinct from the other members of this polymorphic protein family? Metallomics 4:342–349.

Pinharanda, A., S. Martin, S. Barker, J. Davey and C. Jiggins, 2017 The comparative landscape of duplications in Heliconius melpomene and Heliconius cydno. Heredity 118:78–87.

Poelchau, M. F., M.-J. M. Chen, Y.-Y. Lin and C. P. Childers, 2018 Navigating the i5k Workspace@ NAL: a resource for arthropod genomes, pp. 557–577 in Eukaryotic Genomic Databases. Springer.

Pringle, E. G., S. W. Baxter, C. L. Webster, A. Papanicolaou, S. F. Lee and C. D. Jiggins, 2007 Synteny and chromosome evolution in the Lepidoptera: evidence from mapping in Heliconius melpomene. Genetics 177:417–426.

Purać, J., T. V. Nikolić, D. Kojić, A. S. Ćelić, J. J. Plavša, D. P. Blagojević and E. T. Petri, 2019 Identification of a metallothionein gene in honey bee Apis mellifera and its expression profile in response to Cd, Cu and Pb exposure. Molecular ecology 28:731–745.

Ragusa, M. A., A. Nicosia, S. Costa, A. Cuttitta and F. Gianguzza, 2017 Metallothionein gene family in the sea urchin paracentrotus lividus: gene structure, differential expression and phylogenetic analysis. International journal of molecular sciences 18:812.

Rane, R. V., S. L. Pearce, F. Li, C. Coppin, M. Schiffer, J. Shirriffs, C. M. Sgrò, P. C. Griffin, G. Zhang and S. F. Lee, 2019 Genomic changes associated with adaptation to arid environments in cactophilic Drosophila species. BMC genomics 20:52.

Ranz, J. M., F. Casals and A. Ruiz, 2001 How malleable is the eukaryotic genome? Extreme rate of chromosomal rearrangement in the genus Drosophila. Genome research 11:230–239.

Rigby Duncan, K. E., and M. J. Stillman, 2007 Evidence for noncooperative metal binding to the α domain of human metallothionein. The FEBS journal 274:2253–2261.

Robbins, A., D. McRee, M. Williamson, S. Collett, N. Xuong, W. Furey, B. Wang and C. Stout, 1991 Refined crystal structure of Cd, Zn metallothionein at 2.0 Åresolution. Journal of molecular biology 221:1269–1293.

Sims, H. I., G.-W. Chirn and M. T. Marr, 2012 Single nucleotide in the MTF-1 binding site can determine metal-specific transcription activation. Proceedings of the National Academy of Sciences 109:16516–16521.

Smith, G., and A. D. Briscoe, 2015 Molecular evolution and expression of the CRAL_TRIO protein family in insects. Insect biochemistry and molecular biology 62:168–173.

Stewart, N. B., and R. L. Rogers, 2019 Chromosomal rearrangements as a source of new gene formation in Drosophila yakuba. PLoS genetics 15:e1008314.

Thomas, G. W., E. Dohmen, D. S. Hughes, S. C. Murali, M. Poelchau, K. Glastad, C. A. Anstead, N. A. Ayoub, P. Batterham and M. Bellair, 2020 Gene content evolution in the arthropods. Genome biology 21:1–14.

Thurmond, J., J. L. Goodman, V. B. Strelets, H. Attrill, L. S. Gramates, S. J. Marygold, B. B. Matthews, G. Millburn, G. Antonazzo and V. Trovisco, 2018 FlyBase 2.0: the next generation. Nucleic acids research 47:D759–D765.

Vakirlis, N., A.-R. Carvunis and A. McLysaght, 2020 Synteny-based analyses indicate that sequence divergence is not the main source of orphan genes. Elife 9:e53500.

Wang, W.-C., H. Mao, D.-D. Ma and W.-X. Yang, 2014 Characteristics, functions, and applications of metallothionein in aquatic vertebrates. Frontiers in Marine Science 1:34.

Westerman, E. L., N. W. VanKuren, D. Massardo, A. Tenger-Trolander, W. Zhang, R. I. Hill, M. Perry, E. Bayala, K. Barr and N. Chamberlain, 2018 Aristaless controls butterfly wing color variation used in mimicry and mate choice. Current Biology 28:3469–3474. e3464.

Xun, E., Y. Zhang, J. Zhao and J. Guo, 2017 Translocation of heavy metals from soils into floral organs and rewards of Cucurbita pepo: implications for plant reproductive fitness. Ecotoxicology and environmental safety 145:235–243.

Yang, J., and Y. Zhang, 2015 Protein structure and function prediction using I‐TASSER. Current protocols in bioinformatics 52:5.8. 1–5.8. 15.

Yin, C., G. Shen, D. Guo, S. Wang, X. Ma, H. Xiao, J. Liu, Z. Zhang, Y. Liu and Y. Zhang, 2015 InsectBase: a resource for insect genomes and transcriptomes. Nucleic acids research 44:D801–D807.

Zhang, C., P. L. Freddolino and Y. Zhang, 2017 COFACTOR: improved protein function prediction by combining structure, sequence and protein–protein interaction information. Nucleic acids research 45:W291–W299.

Ziller, A., and L. Fraissinet-Tachet, 2018 Metallothionein diversity and distribution in the tree of life: a multifunctional protein. Metallomics 10:1549–1559.

